# SCARECROW is deployed in distinct developmental contexts during rice and maize leaf development

**DOI:** 10.1101/2021.11.29.470347

**Authors:** Thomas. E. Hughes, Jane A. Langdale

## Abstract

The flexible deployment of developmental regulators is an increasingly appreciated aspect of plant development and evolution. The GRAS transcription factor SCARECROW (SCR) regulates the development of the endodermis in Arabidopsis and maize roots, but during leaf development it regulates the development of distinct cell-types; bundle-sheath in Arabidopsis and mesophyll in maize. In rice, SCR is implicated in stomatal patterning, but it is unknown whether this function is additional to a role in inner leaf patterning. Here, we demonstrate that two duplicated *SCR* genes function redundantly in rice. Contrary to previous reports, we show that these genes are necessary for stomatal development, with stomata virtually absent from leaves that are initiated after germination of mutants. The stomatal regulator OsMUTE is down-regulated in *Osscr1;Osscr2* mutants indicating that OsSCR acts early in stomatal development. Notably, *Osscr1;Osscr2* mutants do not exhibit the inner leaf patterning perturbations seen in *Zmscr1;Zmscr1h* mutants and *Zmscr1;Zmscr1h* mutants do not exhibit major perturbations in stomatal patterning. Taken together, these results indicate that SCR was deployed in different developmental contexts after the divergence of rice and maize around 50 million years ago.

**Summary statement:** The transcription factor SCARECROW patterns stomata in rice leaves whereas in maize it predominantly patterns the inner leaf.

## 1. Introduction

The co-ordination of cell patterning is fundamental to multicellularity. In plants, leaf function is underpinned by the correct spatial specification of an array of specialised cell-types. The inner leaf consists of photosynthetic mesophyll cells and a network of vasculature with associated bundle-sheath cells whereas the leaf surfaces comprise epidermal pavement cells, stomatal pores that regulate gas exchange across the leaf surface and hair cells. In grasses, parallel veins within the leaf are flanked by files of stomata in the epidermis (Stebbins & Shah, 1960), requiring co-ordinated development between the inner and outer leaf layers (McKown & Bergmann, 2020).

The GRAS transcription factor SCARECROW (SCR) is one of the best understood plant developmental regulators, having first been identified through its role in regulating cell-type patterning in roots (Laurenzio et al., 1996). In this context, *SCR* is expressed in the initial cells (Laurenzio et al., 1996) that divide asymmetrically to form endodermal and cortical cell-layers around the stele (Dolan et al., 1993). In the absence of SCR, this asymmetric cell division does not occur, resulting in a mutant cell-layer with features of both the endodermis and cortex (Laurenzio et al., 1996). A similar phenotype is seen in maize, where *Zmscr* mutants fail to properly specify the endodermis (Hughes et al., 2019). Furthermore, both Arabidopsis and maize *scr* mutants exhibit a perturbed growth phenotype (Dhondt et al., 2010; Hughes et al., 2019; Hughes & Langdale, 2020). In contrast to these conserved functions, SCR has divergent functions in leaves, patterning bundle-sheath cells around veins in Arabidopsis (Cui et al., 2014; Wysocka-Diller et al., 2000) but regulating mesophyll cell specification and division in maize (Hughes et al., 2019).

Asymmetric cell divisions are a common feature of many plant developmental pathways including stomatal patterning. In eudicots, a meristemoid mother cell divides asymmetrically to give rise to a meristemoid which can then be specified as a guard mother cell (GMC). In monocots, where stomata develop in rows flanking parallel veins, once the stomatal cell file is established cells divide asymmetrically to form a larger interstomatal sister cell and a GMC. In both cases, once established the GMC differentiates and divides symmetrically into the stomatal guard cell pair (reviewed in Conklin et al., 2019; Guo et al., 2021). Consistent with its role in asymmetric cell divisions in root development, it has been suggested that SCR may be required for stomatal patterning in rice (Kamiya et al., 2003; Wu et al., 2019). Unlike in maize, where *ZmSCR* transcripts accumulate primarily in the inner leaf during development (Hughes et al., 2019; Lim et al., 2005) *OsSCR* transcripts accumulate in the epidermis and mark developing stomata (Kamiya et al., 2003). Notably, although duplicate *SCR* genes are now known to be present in both species Kamiya *et al*. (2003) were only aware of one in rice. Given the high level of sequence similarity between the two sequences (CDS 96% sequence similarity), however, it is probable that the in situ hybridisation analysis detected transcripts of both genes (Kamiya et al., 2003). In maize, phenotypic perturbations are only observed in double *Zmscr1;Zmscr1h* mutants (Hughes et al., 2019), whereas in rice *Osscr1* single mutants reportedly showed defects (Wu et al., 2019). Specifically, *Osscr1* but not *Osscr2* mutants exhibited reduced stomatal density, and stomatal patterning in *Osscr1;Osscr2* double mutants was similar to that seen in *Osscr1* single mutants (Wu et al., 2019). Because *SCR* duplicated independently in maize and rice (Hughes et al., 2019) this observation raises the possibility that *OsSCR1* and *OsSCR2* have diverged in function, such that OsSCR1 patterns the epidermis and OsSCR2 patterns the inner leaf layers. This possibility needs further investigation to determine the extent to which SCR function has diverged both within and between rice and maize.

Here we show, contrary to previous reports, that *OsSCR1* and *OsSCR2* genes function redundantly in rice. Double but not single mutants exhibit severe growth and root patterning defects that are similar to those seen in maize and Arabidopsis. However, the rice double mutants do not show the inner leaf patterning defects observed in maize and instead leaves formed post-embryogenesis are almost entirely devoid of stomata, a phenotype far more severe than previously reported. Fittingly, the expression of known stomatal regulators were reduced in double mutants. No such reduction in stomata is found on the abaxial leaf surface in the equivalent maize mutant, although a minor reduction in stomatal density is found on the adaxial surface. Taken together, our results demonstrate that SCR has been recruited into two separate developmental pathways in two closely related monocot species that diverged around 50 million years ago (Vicentini et al., 2008; Wolfe et al., 1989).

## 2. Results

### 2.1 Generation of *Osscr* mutant lines

To assess the role of SCR in rice development four CRISPR guide RNAs (gRNA) were designed (two that targeted *OsSCR1* and two that targeted *OsSCR2*) and cloned into constructs that would enable all relevant combinations of knock-out mutants to be generated and assessed (Fig. 1A,B; Fig. S1). All guides target regions near the 5’ end of the gene sequence such that out-of-frame deletions or insertions lead to complete loss of a functional protein (Fig. 1A). All four guides edited successfully and a number of mutant T0 plants were obtained. Mutant alleles were sequenced and those used for phenotypic characterisation are summarised in Figures 1 & S1. In lines transformed with constructs designed to knock-out both *SCR* genes, it was notable that no T0 plants were identified in which all four alleles were likely to encode non-functional proteins. Instead, all T0 plants screened were found to encode at least one in-frame deletion or insertion and were thus predicted to have at least one functional allele. Given the known perturbed growth phenotypes associated with *scr* mutants in Arabidopsis and maize (Dhondt et al., 2010; Hughes et al., 2019), we hypothesised that complete loss-of-function mutants in rice may not regenerate from tissue culture and/or set seed. Therefore, edited but construct-free T1 plants were identified from two independent lines (both generated using the 17666 construct) in which one allele was predicted to be an in-frame deletion that would likely not substantially alter the protein sequence. As such, plants should grow normally and set T1 seed segregating one in four for a full loss-of-function double mutant. Line 17666-13a contained the alleles *Osscr1-m7/m8;Osscr2-m3/m4* and line 17666-17a contained the alleles *Osscr1-m6/m7;Osscr2-m8/m10* (Fig. 1C). Because the two lines encoded non-identical alleles we deduced that they were independently generated from tissue culture. These two lines, alongside equivalent single mutant lines (Fig. S1E), were prioritised for phenotypic analysis (Fig. 1C).

**Figure 1.**
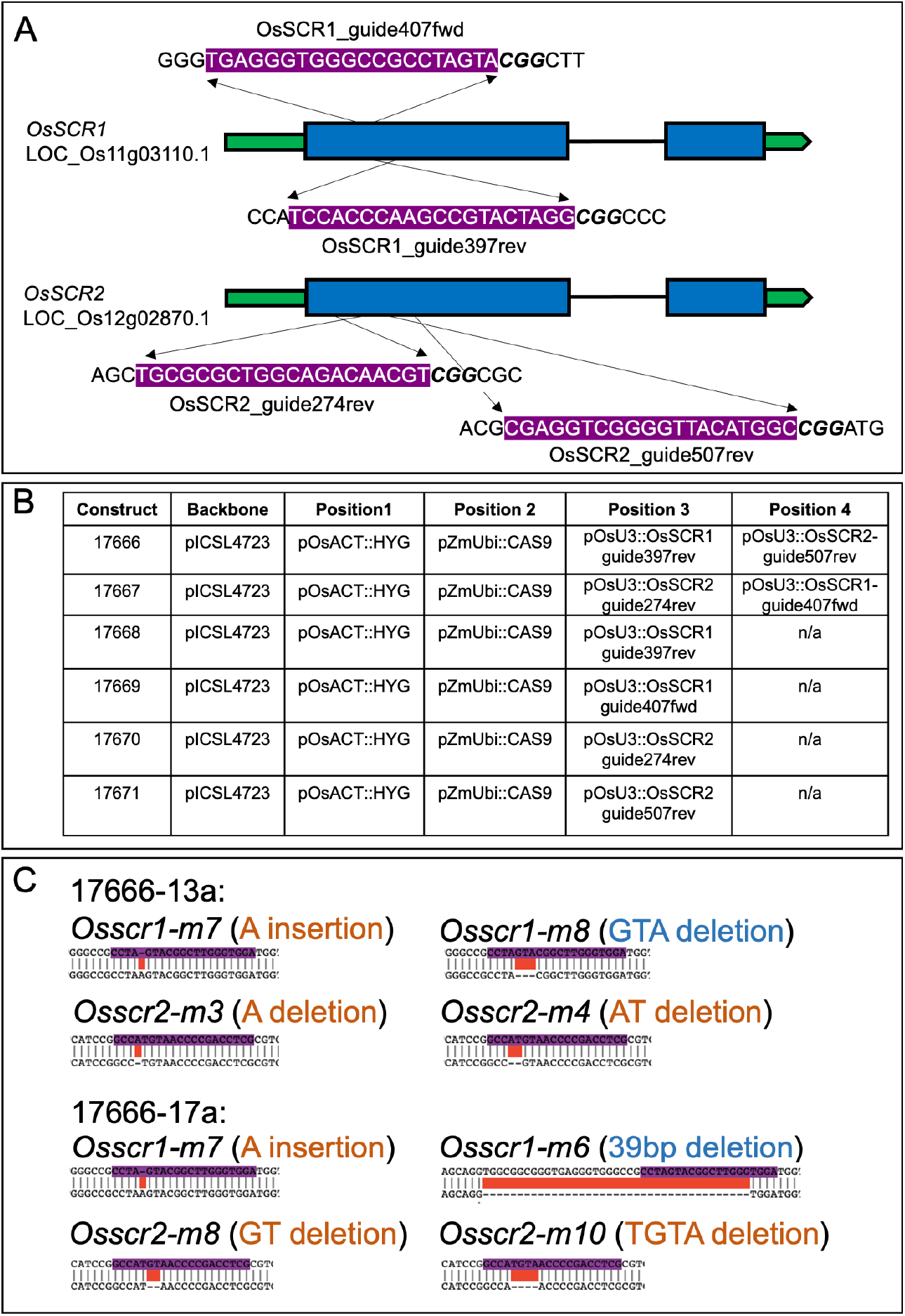
Generation of CRISPR mutants. **A)** Cartoon depiction of guides designed for *OsSCR1* and *OsSCR2* genes. UTR regions are depicted in green, exons in blue, and introns as single black lines. Guide sequences are 5’-3’ and highlighted purple above their rough position either above (forward guides) or below (reverse guides) the gene model. **B)** Table with details of each construct used in the study. **C)** Summary of mutant alleles in two independent double mutants used for phenotypic characterisation. Mutations that lead to a frame-shift of the downstream protein are highlighted in orange, mutations that do not alter the reading frame are highlighted in blue. Sequences are the WT sequence on top and mutant allele sequence beneath.

### 2.2 *Osscr1;Osscr2* double mutants have perturbed growth and root development

Detailed phenotypic analysis was undertaken on the progeny of self-pollinated *Osscr1/+;Osscr2* plants from both independent lines (where + indicates an in-frame edit and thus predicted wild-type protein function). In each case, roughly ¼ of the progeny consistently exhibited a striking growth phenotype whereby plants had very few roots, and the shoots were very small with rolled up leaves (Fig. 2A-F). This phenotype was also observed in other T1 lines that were not taken forward for detailed analysis. In all cases, sequencing confirmed that these plants were homozygous loss-of-function double mutants. The two lines analysed in detail are referred to from here on as *Osscr1-m7;Osscr2-m3* (*1-m7;2-m3* for short) and *Osscr1-m7;Osscr2-m10* (*1- m7;2-m10* for short). Double mutants grew more slowly than wild-type and rarely survived beyond 3-4 weeks after germination. In contrast, both single mutants and plants with at least one in-frame deletion displayed normal growth phenotypes and appeared identical to wild-type (Figure S2), indicating that OsSCR1 and OsSCR2 function redundantly to regulate growth.

**Figure 2.**
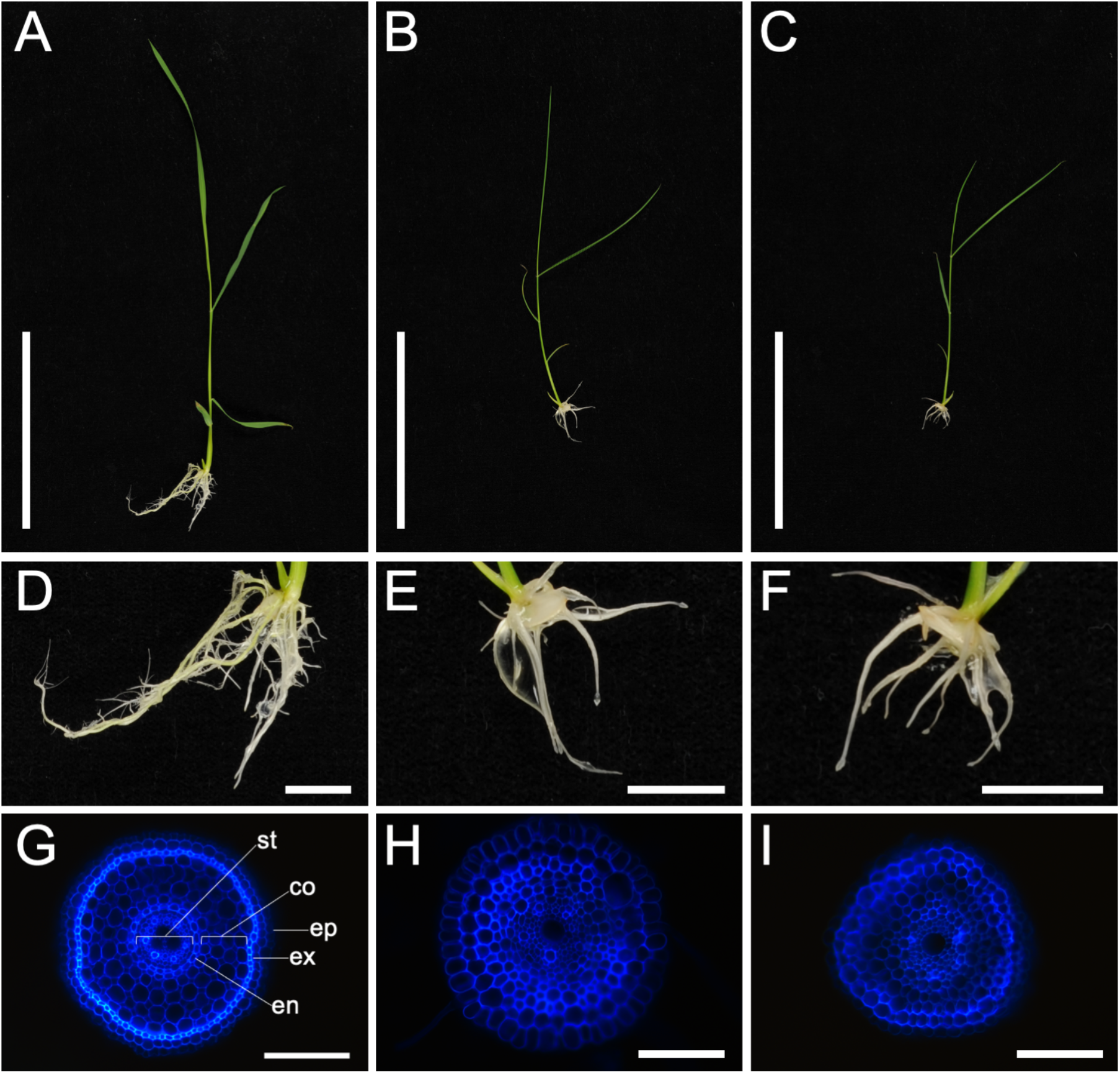
*Osscr1;Osscr2* mutants have perturbed shoot and root development. **A-F)** Photographs of WT (A,D), *Osscr1-m7;Osscr2-m3* (B,E) and *Osscr1-m7;Osscr2- m10* (C,F) plants 14 days after germination. Scalebars in A-C are 10 cm and in D-F are 1 cm. **G-I)** Fresh cross-sections of kitaake WT (G), *Osscr1-m7;Osscr2-m3* (H) and *Osscr1-m7;Osscr2-m10* (I) roots, imaged under ultra-violet illumination. Scalebars in G-I are 100μm. In G the epidermis (ep), exodermis (ex), cortex (co), endodermis (en) and stele (st) are labelled.

It has previously been demonstrated that despite differences in monocot and dicot root development, SCR has a conserved role in endodermal patterning in both maize and Arabidopsis (Hughes et al., 2019; Laurenzio et al., 1996; Lim et al., 2005). To establish whether this role is also conserved in rice we analysed cross-sections of *Osscr1;Osscr2* roots. Because double mutants form fewer and shorter roots than wild-type (Fig. 2A-F), we compared unbranched roots with similar diameters and obtained cross-sections from the maturation zone above the root tips (Fig. 2G-I). *Osscr1;Osscr2* roots displayed a severely perturbed phenotype. Specifically, cell layers were disorganised throughout with no obvious endodermal layer separating the vasculature from the cortex, and no obvious exodermal layer separating the cortex from the epidermis (Fig. 2G-I). This phenotype strongly resembles that found in maize *Zmscr1;Zmscr1h* roots, indicating that the role of SCR in root patterning is conserved in rice, maize and Arabidopsis.

### 2.3 *Osscr1;Osscr2* double mutants show no obvious inner leaf patterning defects

In maize, *Z*mSCR1 and ZmSCR1h function redundantly to regulate mesophyll specification and cell division in the inner leaf (Hughes et al., 2019). To assess whether this function is conserved in rice, which has many more mesophyll cells positioned between veins than maize, we examined cross-sections of *Osscr1;Osscr2* mutant leaves (Fig. 3A-F). Despite being significantly narrower (Fig. 3G), and more rolled (Fig. 3B,C) double mutant leaves did not exhibit any patterning defects in the inner leaf. Although vein density was significantly increased (Fig. 3H) and inter-veinal distance was significantly reduced (Fig. 3I) these effects are likely to be caused by altered leaf width and/or cell size rather than by any direct cell-patterning defects. In support of this suggestion, traits that are characteristic of patterning defects in *Zmscr1;Zmscr1h* mutants were absent. That is, there was no reduction in the number of mesophyll cells separating veins (Fig. 3J), no ectopic bundle-sheath or sclerenchyma cells, and no fused vascular bundles (Fig. 3A-C). In summary, no evidence was found to support a role for *OsSCR1* or *OsSCR2* in inner leaf patterning in rice, indicating that in this context SCR function has diverged between rice and maize.

**Figure 3.**
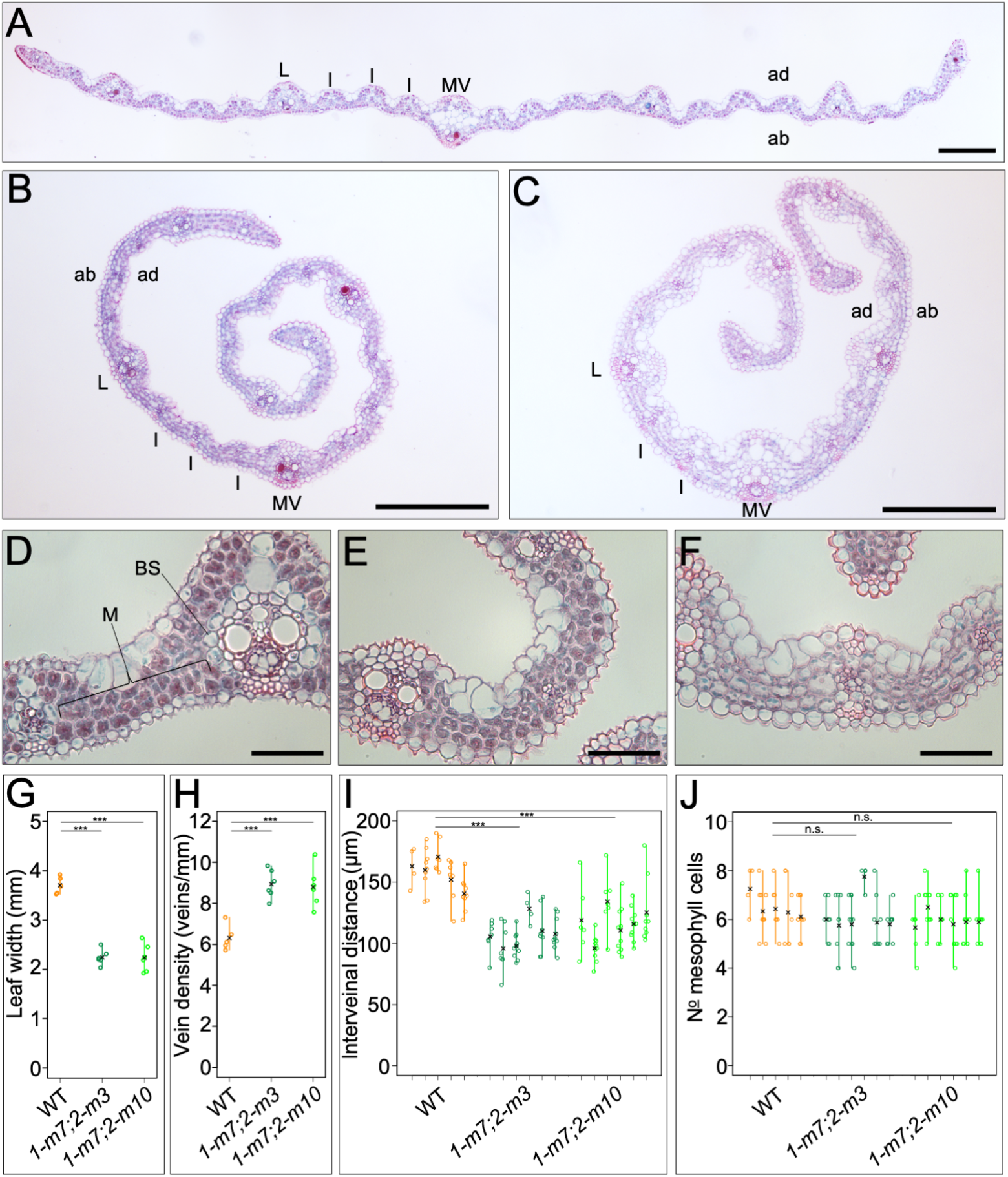
Inner leaf cell patterning is unperturbed in *Osscr1;Osscr2* mutants. **A-F)** Cross-sections of WT (A,D), *Osscr1-m7;Osscr2-m3* (B,E) and *Osscr1-m7;Osscr2- m10* (C,F) leaves. Leaves were sampled from the mid-point of fully-expanded leaf 5, 18 days after germination. In A-C the mid-vein (MV), examples of lateral (L) and intermediate (I) veins, and the abaxial (ab) and adaxial (ad) surfaces are indicated, in D mesophyll (M) and bundle-sheath (BS) cells are indicated. Scalebars are 100μm in A-C and 50μm in D-F. **G-J)** Quantification of leaf width (G), vein density (H), inter- veinal distance (I) and the number of mesophyll cells separating veins (J). The means for each plant are indicated by a black cross. In I and J each line represents an individual leaf, and each circle represents one inter-veinal region. Statistical significance between each genotype was assessed using one-way ANOVA and TukeyHSD posthoc tests: n.s. *P*>0.05; \**P*≤0.05; \*\**P*≤0.01; \*\*\**P*≤0.001.

### 2.4 OsSCR1 and OsSCR2 function redundantly to specify stomata

Given our finding that OsSCR1 and OsSCR2 function redundantly to regulate plant growth, we found the reported stomatal defects in *Osscr1* single mutants perplexing (Wu et al., 2019). We therefore sought to better understand the role of both genes in stomatal development (Figure 4). In wild-type rice, three to four leaf primordia are initiated and at least partially patterned during embryogenesis (Itoh et al., 2005). To examine stomatal patterning in leaves formed both during embryogenesis and post-germination we therefore quantified stomatal density on leaves 3, 4 and 5, using resin impressions of the leaf surface (Fig. 4A-I). Surprisingly, given previously published results, stomatal density was not reduced on the abaxial surface of any leaf in either *Osscr1* or *Osscr2* single mutants (Fig. 4J-L). In contrast, abaxial stomatal density was reduced to around 60% of wild-type in leaf 3 of double *Osscr1;Osscr2* mutants (Fig. 4A,D,G,J), and stomatal rows were not as clearly defined, with instances of clustering (Fig 4. D,G). These phenotypes were even more pronounced in leaves 4 and 5, with stomata rarely observed in leaf 4 (Fig. 4B,E,H,K) and never observed in leaf 5 of the *Osscr1-m7;Osscr2-m10* mutant (Fig. 4F,K). A few stomata were observed in leaf 5 of the *Osscr1-m7;Osscr2-m3* mutant, but only in two of five individual plants examined (Fig. 4C,F,I,L). When stomata were present, they were in patches and in poorly defined rows (Fig. 4E,H). The adaxial surface of leaf 5 was also devoid of stomata in both double mutants (Fig. S3A-C). To confirm that these results were not a technical artefact of resin impressions, double mutant leaves were also examined directly under a scanning electron microscope. The images obtained confirmed the virtual absence of stomata on both surfaces of leaf 5 (Fig. S3D-G). No obvious aborted stomata or guard mother cells (GMCs) were seen in either double mutant, although assessing epidermal patterning of rice leaves is made challenging by the presence of wax and silica on the leaf surface (Figure 4, Figure S3). In summary, OsSCR1 and OsSCR2 redundantly regulate stomatal development, particularly in leaves that are initiated post-embryogenesis.

**Figure 4.**
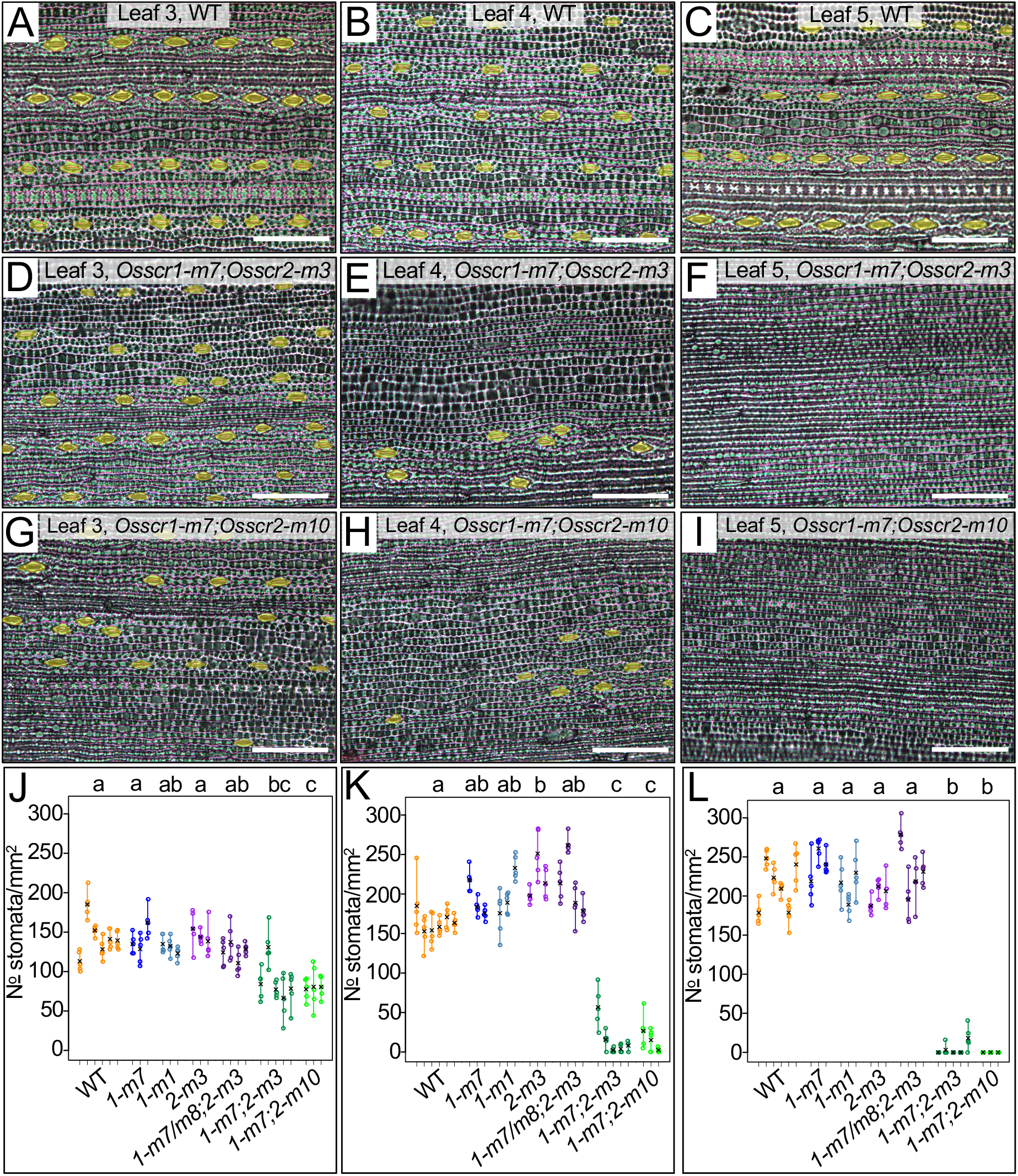
*Osscr1;Osscr2* leaves have almost no stomata. **A-I)** Abaxial impressions of WT (A-C), *Osscr1-m7;Osscr2-m3* (*1-m7;2-m3*) (D-F) and *Osscr1-m7;Osscr2-m10* (*1-m7;2-m10*) (G-I) leaves 3 (A,D,G), 4 (B,E,H) and 5 (C,F,I). Stomata are false coloured yellow. Leaves were sampled at the mid-point along the proximal-distal axis either 14 (leaves 3 and 4) or 18 days (leaf 5) after germination. Scalebars are 100μm. **J-L)** Quantification of stomatal density of leaf 3 (J), leaf 4 (K) and leaf 5 (L) for WT, *Osscr1-m7* (*1-m7*), *Osscr1-m1* (*1-m1*), *Osscr2-m3* (*2-m3*), *Osscr1-m7/m8;Osscr2-m3* (*1-m7/m8;2-m3*) (where *1-m8* is an in-frame deletion and thus functionally these are *Osscr2* single mutants) and double mutants. Different colours depict the different genotypes beneath each group. Each line represents an individual leaf, and each circle an image quantified for each leaf. The means for each plant are indicated by a black cross. Letters at the top of each plot indicate statistically different groups (*P* < 0.05, one-way ANOVA and TukeyHSD).

### 2.5 OsSCR functions upstream of OsMUTE during stomatal development

A number of genes that regulate stomatal specification and differentiation have been identified in monocots, largely due to having conserved roles with Arabidopsis orthologs. For example, in rice and Brachypodium, MUTE regulates GMC differentiation and subsidiary cell recruitment and FAMA regulates final stomatal patterning (Liu et al., 2009; Raissig et al., 2017; Wu et al., 2019). SPEECHLESS (SPCH) has also been implicated in controlling entry to the stomatal lineage in monocots (Raissig et al., 2016; Wu et al., 2019), but in line with previous studies in rice we were unable to reliably detect SPCH expression at quantifiable levels (Liu et al., 2009). Therefore, to position SCR in this pathway we quantified *OsMUTE* and *OsFAMA* transcripts in developing *Osscr1;Osscr2* mutant leaves 6 days after sowing (Figure 5). Given that the perturbed growth phenotype in *Osscr1;Osscr2* mutants may represent a developmental delay relative to wild-type, we first quantified *ROC5* gene transcript levels. *ROC5* has been previously shown to mark the developing epidermis in rice (Ito et al., 2003). Although *ROC5* levels appeared to be slightly reduced in the *Osscr1-m7;Osscr2-m3* line, this was not significant, and there was no consistent reduction in the *Osscr1-m7;Osscr2-m10* line (Fig. 5A). Therefore, we concluded that the relative amount of developing epidermal tissue is similar in both genotypes. In contrast, *OsMUTE* levels were drastically reduced in both *Osscr1;Osscr2* lines (Fig. 5A). There was a trend for reduced *OsFAMA* levels, but to a lesser extent and in a less consistent manner than *OsMUTE* (Fig. 5A). These results were confirmed using WT tissue harvested 4 days after sowing, when shoots were similarly sized to mutant seedlings harvested 6 days after sowing. In this comparison, *ROC5* levels were slightly higher (though not statistically so) than in the mutant, but *OsMUTE* and *OsFAMA* transcript levels in the mutant showed the same pattern as seen in the previous comparison, with *OsMUTE* in particular strongly down-regulated (Fig. 5B). Taken together, these data indicate that OsSCR1 and OsSCR2 function upstream of OsMUTE and OsFAMA in stomatal development, and thus that SCR may regulate entry into the stomatal specification pathway.

**Figure 5.**
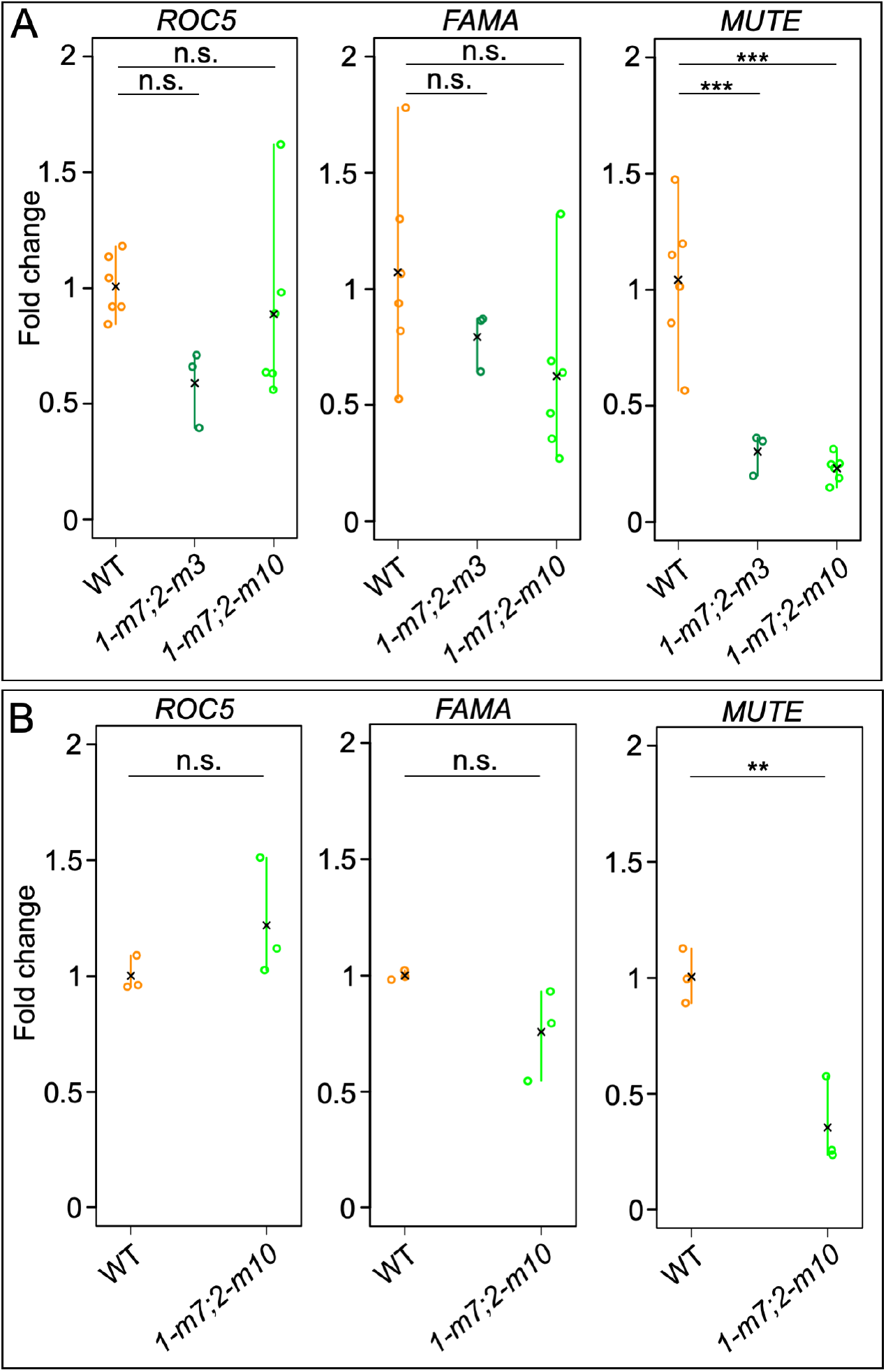
*OsMUTE* is strongly down-regulated in *Osscr1;Osscr2* mutants. **A-B)** Quantitative RT-PCR analysis of *OsROC5, OsFAMA* and *OsMUTE* levels in WT, *Osscr1-m7;Osscr2-m3* (*1-m7;2-m3*) and *Osscr1-m7;Osscr2-m10* (*1-m7;2-m10*) plants. In A both kitaake WT and mutant seedlings were harvested 6 days after germination, whereas in B kitaake WT was harvested 4 days after germination and compared to the same mutant samples in A. In each plot means are indicated by a black cross. Statistical significance between each genotype was assessed using one- way ANOVA and TukeyHSD posthoc tests: n.s. *P*>0.05; \**P*≤0.05; \*\**P*≤0.01; \*\*\**P*≤0.001.

### 2.6 Stomatal density is only reduced on the adaxial surface of *Zmscr1;Zmscr1h* mutants

To assess whether a role in stomatal development is specific to rice or shared with other monocots, we assessed stomatal patterning in leaves 4 to 7 of two independent *Zmscr1;Zmscr1h* mutants (Fig. 6) (Hughes et al., 2019). Maize initiates up to 5 leaves during embryogenesis (W.-Y. Liu et al., 2013) and thus leaves 4 to 7 encompass the same developmental trajectory from embryonic to non-embryonic leaves as rice leaves 3 to 5. Whereas leaf 5 of *Osscr1;Osscr2* mutants formed very few stomata on either the abaxial or adaxial surface, leaf 7 of *Zmscr1;Zmscr1h* mutants showed no reduction in stomatal density on the abaxial surface (Fig 6. A-H, Q-T). There was a decrease in stomatal density on the adaxial surface (Fig 6. I-P, U-X), however, the reduction was not as great as seen in rice and there was no difference between embryonic and non-embryonic leaves. Given that both *ZmSCR1* and *ZmSCR1h* are expressed in the inner layers of the maize leaf (Hughes et al., 2019) rather than in developing stomata, the reduction in adaxial stomatal density in double mutants is likely an indirect consequence of loss of SCR function, although it is possible that some aspect of a stomatal patterning function is retained in maize. Whether an indirect or direct effect, however, the stomatal phenotype in maize double mutants is so distinct from that exhibited in rice double mutants that there is a high degree of divergence in SCR function between these species.

**Figure 6.**
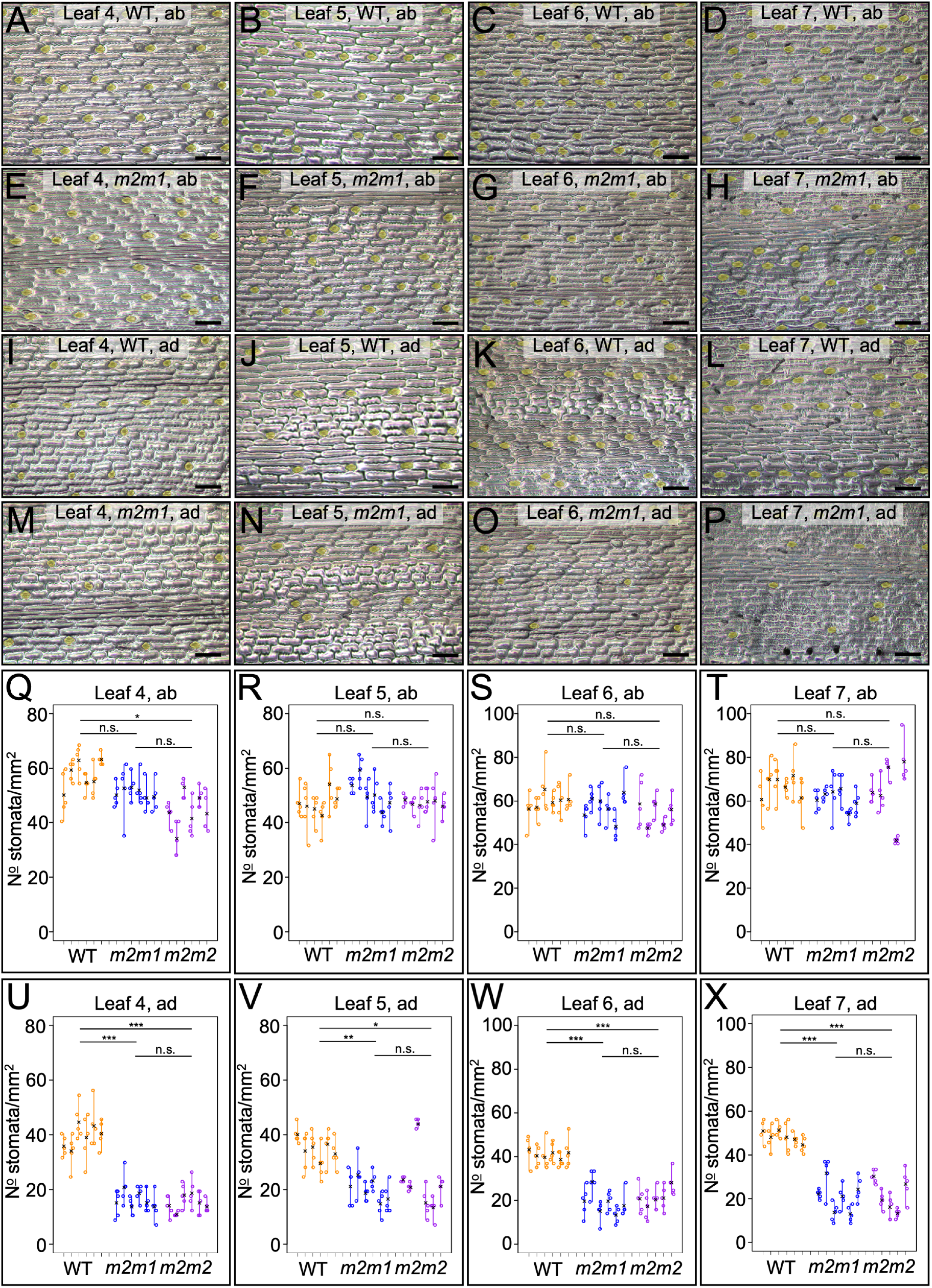
Adaxial but not abaxial stomatal density is reduced in *Zmscr1;Zmscr1h* mutants. **A-P)** Epidermal impressions of abaxial (A-H) or adaxial (I-P) maize leaves from either segregating WT (A-D,I-L) or *Zmscr1-m2;Zmscr1-m1* (*m2m1*) mutants (E- H,M-P), for leaves 4 (A,E,I,M), 5 (B,F,J,N), 6 (C,G,K,O) and 7 (D,H,L,P). Scalebars are 100μm. Stomata are false coloured yellow. **Q-X)** Stomatal density quantification for both abaxial (Q-T) and adaxial (U-X) leaf surfaces for leaves 4 (Q,U), 5 (R,V), 6 (S,W) and 7 (T,X). In each plot in (Q-X) each line is an individual plant, and each datapoint an image quantified for that plant. Different colours depict the segregating m2m1 WT line (orange), *m2m1* mutant (blue) and *m2m2* mutant (purple). Means are depicted by a black cross. Statistical significance between each genotype was assessed using one-way ANOVA and TukeyHSD posthoc tests: n.s. *P*>0.05; \**P*≤0.05; \*\**P*≤0.01; \*\*\**P*≤0.001

## 3. Discussion

It is increasingly appreciated that the same gene or regulatory module can be co-opted to regulate distinct developmental pathways both within and between species. In this study we have demonstrated that in rice, duplicate *SCR* genes redundantly regulate growth and root development in a manner that is shared with orthologous genes in Arabidopsis and maize (Fig. 2). In the context of leaf development, however, the rice genes have a novel role. Gene function is necessary for stomatal development, with loss of function mutants virtually devoid of stomata on the leaf surface (Fig. 4). We position OsSCR1 and OsSCR2 upstream of the known stomatal regulators OsMUTE and OsFAMA (Fig. 5). The stomatal patterning perturbations in rice mutants are not exhibited in equivalent maize mutants (Fig. 6), and the inner leaf patterning perturbations described in maize mutants are not apparent in rice (Fig. 3). Taken together, these results demonstrate that in the context of leaf development, SCR has distinct roles in two closely related monocot species.

Previous reports have suggested that OsSCR1 plays a more important role in stomatal development than OsSCR2 (Wu et al., 2019). Wu et al. (2019) found that stomatal density was reduced to around 50% that of wild-type in leaf 5 of *Osscr1* single mutants, and that the *Osscr1;Osscr2* mutant phenotype was only slightly more severe. This stands in marked contrast to our results, where no decrease was seen in the single mutant whereas stomata were virtually absent in leaf 5 of the double mutant. It is not clear why these differences have arisen. Although an environmental contribution cannot be ruled out, our results clearly demonstrate that contrary to previous reports, OsSCR1 and OsSCR2 function redundantly in stomatal development.

The regulation of stomatal development in eudicots is well characterised (reviewed in Simmons & Bergmann, 2016), but it is only recently that equivalent regulators have been identified in monocots. Monocot grasses such as rice, maize and *Brachypodium* have complex stomata that develop subsidiary cells in addition to guard cell pairs (Stebbins & Shah, 1960). In *Brachypodium* and rice, stomatal cell files are initiated by a complex of three bHLH transcription factors (INDUCER OF CBF EXPRESSION 1 (ICE1), SPEECHLESS1 (SPCH1) and SPCH2) (Raissig et al., 2016; Wu et al., 2019).

Within these cell files, asymmetric cell divisions lead to the formation of the GMCs. In *Brachypodium*, rice and maize, MUTE orthologs function both cell-autonomously to regulate the symmetrical division of the GMC to form guard cells and non-cell autonomously to recruit subsidiary cells from adjacent cell files (Raissig et al., 2017; H. Wang et al., 2019; Wu et al., 2019). Our analysis supports a role for OsSCR1 and OsSCR2 upstream of MUTE because *OsMUTE* transcript levels were severely downregulated in *Osscr1;Osscr2* mutants. Given the absence of GMCs on leaf 5 of *Osscr1;Osscr2* mutants, the expression of *OsSCR* in developing stomata (Kamiya et al., 2003), and the role of SCR in promoting asymmetric cell divisions in Arabidopsis roots (Laurenzio et al., 1996), we hypothesise that in rice (but not maize) SCR functions in stomatal cell files to regulate the asymmetric cell divisions that give rise to GMCs.

In rice, stomata are arranged in files flanking both sides of underlying veins and it has been hypothesised that a positional signal emanating from the veins acts to initiate development of those files (McKown & Bergmann, 2020; Nunes et al., 2020). In Arabidopsis roots, the SHORTROOT (SHR) protein acts as a positional signal, moving from the vasculature to the surrounding cell-layer where it is bound by SCR. Activation of downstream targets by SHR/SCR then leads to an asymmetric cell division and the formation of the endodermis and cortex (Cui et al., 2007). The previous finding that *SCR* genes are expressed in developing stomata of rice leaves (Kamiya et al., 2003), and our validation of a redundant role for these genes in regulating stomatal development, suggests that in rice SHR may signal from the veins to the overlying epidermal regions. Signalling from the inner leaf to the epidermis has recently been demonstrated in Arabidopsis, where the light responsive transcription factor HY5 induces STOMAGEN expression in the mesophyll, which then induces changes in stomatal patterning in the epidermis (Wang et al., 2021). Rice encodes two *SHR* orthologs, one of which is also expressed in developing stomata, albeit less obviously than seen for *SCR* (Kamiya et al., 2003). The second *OsSHR* gene is expressed at low levels in developing veins and when it is ectopically expressed in the bundle sheath cell layer surrounding the vein, additional stomatal files are initiated (Schuler et al., 2018). Given this finding, although there are several unresolved questions (such as how SHR can act as a vein-derived positional signal when one ortholog appears to be expressed in stomata), it is plausible that SCR acts cell-autonomously in stomatal cell files, interpreting a SHR signal from within the rice leaf to trigger the asymmetric cell divisions that generate GMCs.

In maize, *SCR* is not expressed in developing stomata (Hughes et al., 2019; Lim et al., 2005), and the phenotypes of the rice and maize mutants are highly diverged both in the inner leaf and the epidermis. Maize utilises the C_4_ photosynthetic pathway, which is underpinned by Kranz anatomy, whereby vein density is higher than in species such as rice that utilise the ancestral C_3_ pathway. Notably, the higher vein density is not accompanied by higher stomatal density, and as such some veins are not flanked by stomatal files. Given this disconnect it is tempting to speculate that stomatal patterning in C_4_ monocots does not need to be as tightly integrated with venation patterning and thus that SCR is not necessary for stomatal development in maize. Of course, with only one C_3_ and one C_4_ monocot species characterised so far it is not possible to infer which function is ancestral, nor to assess whether the divergent functions reflect specific differences between maize and rice or general differences between C_4_ and C_3_ species. If a role in stomatal patterning is linked to C_3_ leaf development it is likely to be confined to the monocots, because in the C_3_ eudicot Arabidopsis stomatal patterning is unaffected in *scr* mutants (Dhondt et al., 2010). Taken together, these results indicate that in the context of leaf development, SCR function has been recruited into three largely distinct roles in maize, rice and Arabidopsis, namely patterning of mesophyll, stomatal and bundle-sheath cells, respectively.

## 4. Materials and methods

### 4.1 Plant growth

Seed of *Oryza sativa spp japonica* cv Kitaake (referred to as WT or wild-type throughout) were dehulled and sterilised by treatment with 70% ethanol (v/v) for 2 min followed by 25% sodium hypochlorite solution with a drop of tween-20 for 15 min. Sterilised seed were rinsed five times with sterile deionised water and then placed on ½ MS media (2.15g/L Murashige & Skoog salts and vitamins, 0.5g/L MES, 4g/L Phytagel, pH 5.8) in an incubator (Panasonic, MLR-352) with 16 hour photoperiod and 30°C/25°C day-time/night-time temperature. After 7 days, seedlings were transferred to 50ml falcon tubes and watered with ¼ MS solution (1.07g/L Murashige & Skoog salts and vitamins, pH 5.8). Falcon tubes were covered with clingfilm to maintain higher humidity until seedlings emerged out of the tube. Phenotyping was undertaken on plants growing in falcon tubes because *Osscr1;Osscr2* mutants did not survive beyond 3-4 weeks after germination. Plants for seed propagation were transferred into 7.5cm pots filled with clay granules (Profile, Porous Ceramic Topdressing and Construction Material) and placed in a controlled environment chamber with the same conditions as the incubator, and light intensity 250-300μmol photons m^-2^ s^-1^. Trays of 15 plants were covered for around 1 week after transfer with a transparent bag to increase humidity. Plants were watered 3 times a week with a fertiliser solution (1.34g/L Everris Peters Excel Cal-Mag Grower N.P.K. 15-5-15, pH 5-6), with 0.5g/L chelated iron added to the solution on alternate weeks until flowering. Maize seedlings were grown as described previously (Hughes et al., 2019).

### 4.2 CRISPR construct design and cloning

The rice SCR orthologs were obtained from a previously published phylogeny (Hughes et al., 2019). Sequences for each gene were obtained from phytozome V12 (https://phytozome-next.jgi.doe.gov) and guides targeting the first exon of each gene were designed using CRISPOR (Concordet & Haeussler, 2018). Guide oligonucleotides were synthesised with 4bp golden gate sequences and Esp3I restriction sites added to facilitate cloning into a golden gate module. All golden gate reactions were carried out using a standard one-tube reaction as described previously (Engler et al., 2008). Complementary oligonucleotides were mixed in a 1:1 ratio and heated to 99°C before being left to cool for an hour at room temperature and anneal.

Annealed guides were diluted 200-fold for cloning into either module EC15768 (position 3, reverse) or EC15769 (position 4, reverse) (Fig S1A). The resultant level 1 modules contained each guide in a full RNA scaffold sequence, driven by the OsU3 promoter (Fig. S1A). Promoter-guide modules were assembled into final level 2 transformation constructs in the pICSL4723 backbone with hygromycin resistance and CAS9 modules (Fig. S1A).

### 4.3 Rice transformation and tissue culture

Constructs were transformed into *Agrobacterium tumefaciens* strain EHA105 and then used to transform Kitaake rice using a modified transformation protocol (Toki et al., 2006), that can be downloaded here (https://langdalelab.files.wordpress.com/2018/06/kitaake-rice-transformation.pdf).

### 4.4 Genotyping

Initial screening of T0 plants was undertaken with genomic DNA extracted using a modified sodium dodecyl sulfate (SDS) 96-well plate method (Hughes et al., 2019), whereby leaf tissue was frozen in liquid nitrogen and homogenised prior to the addition of extraction buffer. Hygromycin primers (Fig. S1B) were used to screen for successfully transformed T0 plants. Primers were designed to specifically amplify the first exon of either *OsSCR1* or *OsSCR2* (Fig. S1B), which worked efficiently when template genomic DNA was extracted using a previously published CTAB method (Hughes et al., 2019). The resultant amplicon was digested with a restriction enzyme predicted to cut at the intact guide site (Fig S1C). If undigested (and thus edited), the amplicon was cloned into pJET (CloneJET, Thermofisher) and colonies sequenced by Sanger sequencing until both allele sequences were known. Seed were collected from T0 plants of interest and T1 progeny grown to identify mutated plants that lacked the construct. This was achieved using the same hygromycin primers alongside a pair of primers amplifying the rice ubiquitin gene, to ensure that the failure to amplify hygromycin was not due to low DNA quality. Specific genotyping assays were designed to distinguish pairs of alleles in two independent T1 *Osscr1;Osscr2* mutants (Fig. S1D). All PCR reactions were undertaken with GoTaq DNA polymerase (Promega) with cycling conditions 95°C for 5 min; 35 cycles of 95°C for 30 s, 57-61°C for 30 s and 72°C for 60-90 s; and 72°C for 5 min. 1M Betaine (Sigma Aldrich) was added to all amplifications of both *SCR* genes due to high-GC content. Restriction digestions were undertaken directly on the amplified PCR product, with 3μl of a 10μl PCR reaction used in a 10μl digestion at the recommended digestion temperature overnight.

### 4.5 Histology

Seminal roots were cut in the maturation zone of both Kitaake WT and *Osscr1;Osscr2* mutants 15 days after sowing, and positioned in 3% agarose blocks. Roots were sectioned on a Leica VT1200S vibratome at 60μm thickness and the resultant sections floated on slides in deionised water. Sections were imaged using a Leica DMRB microscope and ultraviolet illumination with a DFC7000T camera and Leica LASX image analysis software.

Inner leaf phenotyping was undertaken on leaf segments cut at the mid-point of fully expanded leaf 5 at 18 days after sowing and fixed by submersion in 3:1 ethanol: acetic acid for 30 minutes followed by storage in 70% ethanol. Leaf tissue was wax infiltrated using a Tissue-Tek VIP machine (Sakura, www.sakura.eu) using the protocol published previously (Hughes et al., 2019), and the resultant wax blocks sectioned at 10μm. Sections were cleared using Histoclear (10 min, x2), followed by submersion in 100% and 70% ethanol (both 2 min x2). Sections were stained using safranin-O (1% in 50% ethanol) for 90 mins, rinsed in 70% ethanol (2 min, x2), and counter-stained with fast-green (0.03% in 95% ethanol) for 3 seconds per slide. Finally, slides were rinsed in 100% ethanol (2 min, x2), then 100% histoclear (5 min) and mounted using a drop of DPX mounting medium. Images were obtained using brightfield illumination on the same microscope and software described above.

Stomatal impressions were taken using dental resin (Perfection Plus, Impress Plus Medium Body Fast Set) applied to the mid-point of leaves 3, 4 and 5 on either the abaxial or adaxial surface. Once applied, resin was left to dry for 5 minutes and then the leaf removed. Clear nail varnish (Rimmel) was applied to the dental resin impressions and left to dry for at least 5 minutes, before being peeled off and floated on deionised water which was blotted off to dry impressions to the slide. Sections were imaged using phase-contrast illumination on the same microscope described above. For quantification, five 10x images were taken from random parts of each peel, and a higher magnification 20x image taken for presentation. Stomata were counted and divided by the area of peel to calculate stomatal density. For scanning electron microscopy rice leaves were attached directly to a stub without any pre-treatment and imaged directly on a scanning electron microscope (JEOL JSM5510, 15kV).

### 4.6 Quantitative RT-PCR analysis

Wild type and mutant seed were sterilised and plated on ½ MS media as described above. After either 4 (WT) or 6 (WT and mutant) days, whole shoots from each plant were removed and the length of each measured before freezing in liquid nitrogen. Total RNA was extracted using the Qiagen RNeasy kit and DNA contamination removed using Turbo DNase. 2μg of RNA was used for cDNA synthesis, and cDNA quality checked with ubiquitin primers that amplify distinct products from genomic DNA and cDNA.

Quantitative RT-PCR analysis was undertaken using SYBR-green with cycle conditions 95°C for 10 min, then 40 cycles of 95°C for 15 s and 60°C for 1 min. Melt-curves were obtained between 60°C and 95°C to establish that a single product was amplified for each primer pair. Primers for two housekeeping genes (OsACTIN and OsUBQ5) were obtained from previously published work (Jain et al., 2006; P. Wang et al., 2017), and primers for *OsMUTE* (LOC_Os05g51820), *OsFAMA* (LOC_Os05g50900) and *OsROC5* (LOC_Os02g45250) were designed in the CDS of each gene using Primer3Plus. RT-PCR was undertaken to confirm primers amplified an amplicon of the correct size, and primer efficiency confirmed to be >80% using the qPCR miner algorithm from a test qRT-PCR run (Zhao & Fernald, 2005). Three technical replicates were obtained for each sample and confirmed to have Ct values with a range <∼0.5 once outliers were removed. All comparisons were run on the same plate alongside water controls, and as such Kitaake WT samples were repeated alongside both independent mutant backgrounds. Ct values were calculated using the qPCR miner algorithm (Zhao & Fernald, 2005), and fold-change values using the 2−^ΔΔCT^ method (Livak & Schmittgen, 2001). The overall average wild-type across all samples was used to compare each individual wild-type to indicate the range of the wild-type data. Mutant samples were then compared to the same overall wild-type average, and as such values of less than 1 indicate a relative reduction compared to wild-type.

## Author contributions

T.E.H and J.A.L conceived and designed the experiments. T.E.H carried out the experiments and analysed the data. T.E.H and J.A.L wrote the manuscript.

## Acknowledgements

The authors thank Robert Caine and Michael Raissig for advice on stomatal phenotyping; John Baker for plant photography; Roxaana Clayton, Julie Bull and Lizzie Jamison for technical support; Niloufer Irani for help with scanning electron microscopy; Sophie Johnson, Chiara Perico, Daniela Vlad, Sovanna Tan, Julia Lambret-Frotte and Maricris Zaidem for discussion throughout the experimental work and during manuscript preparation; and Laura Moody and Maddy Seale for providing feedback on the finished manuscript.

---

**Figure S1.**
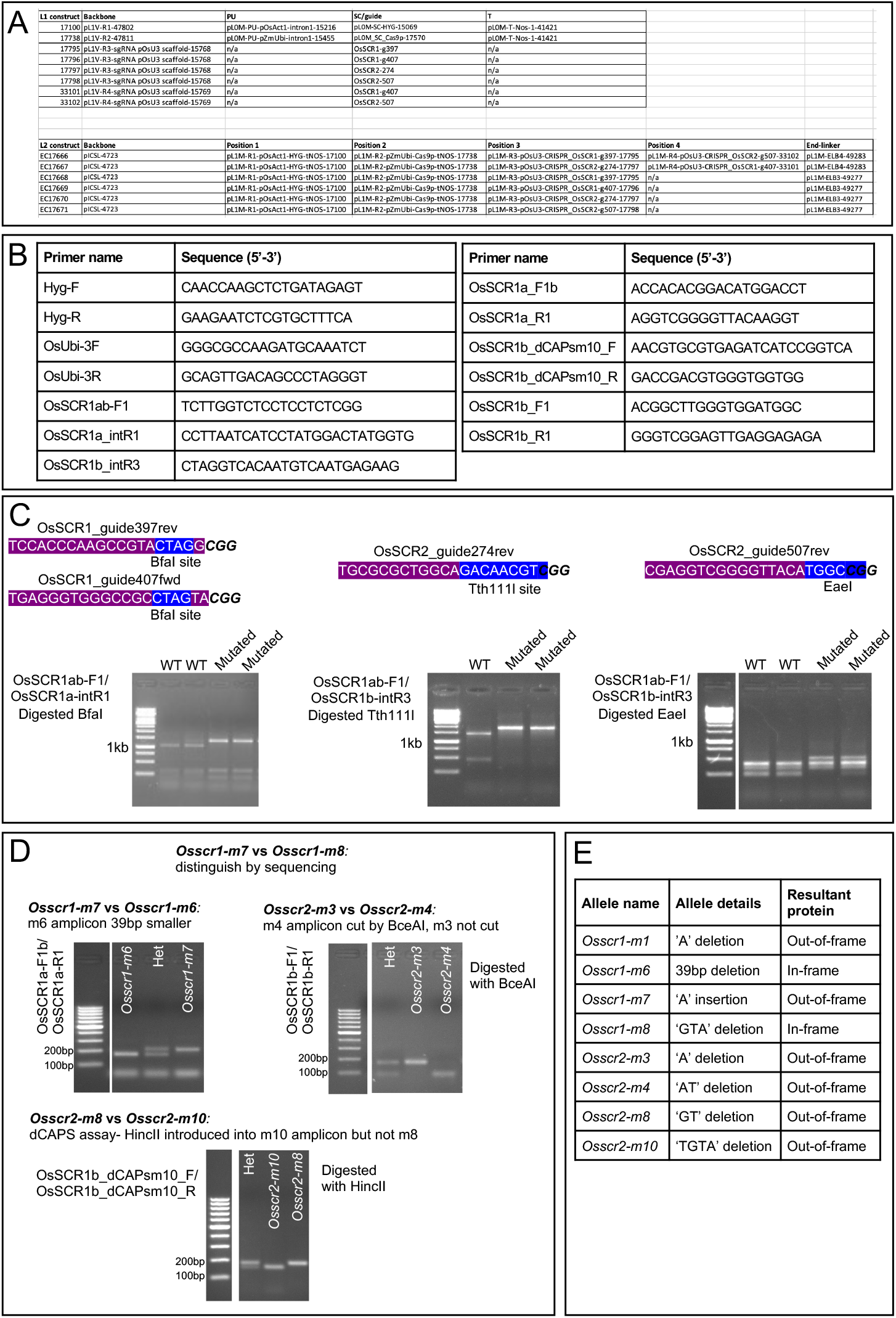
Generation of CRISPR mutants. **A)** Golden gate modules and cloning used in this study. **B)** Genotyping primers used in this study. **C)** Restriction digest assays used to identify mutant alleles for each CRISPR guide. Guide sequence is depicted with purple highlighting and white lettering, the protospacer adjacent motif (PAM) sequence is depicted in black italic text, and the restriction enzyme site is depicted with blue highlighting. **D)** Distinguishing pairs of known alleles identified in Fig. 1C using different restriction enzyme assays. **E)** Summary of all alleles used for phenotyping in this study.

**Figure S2.**
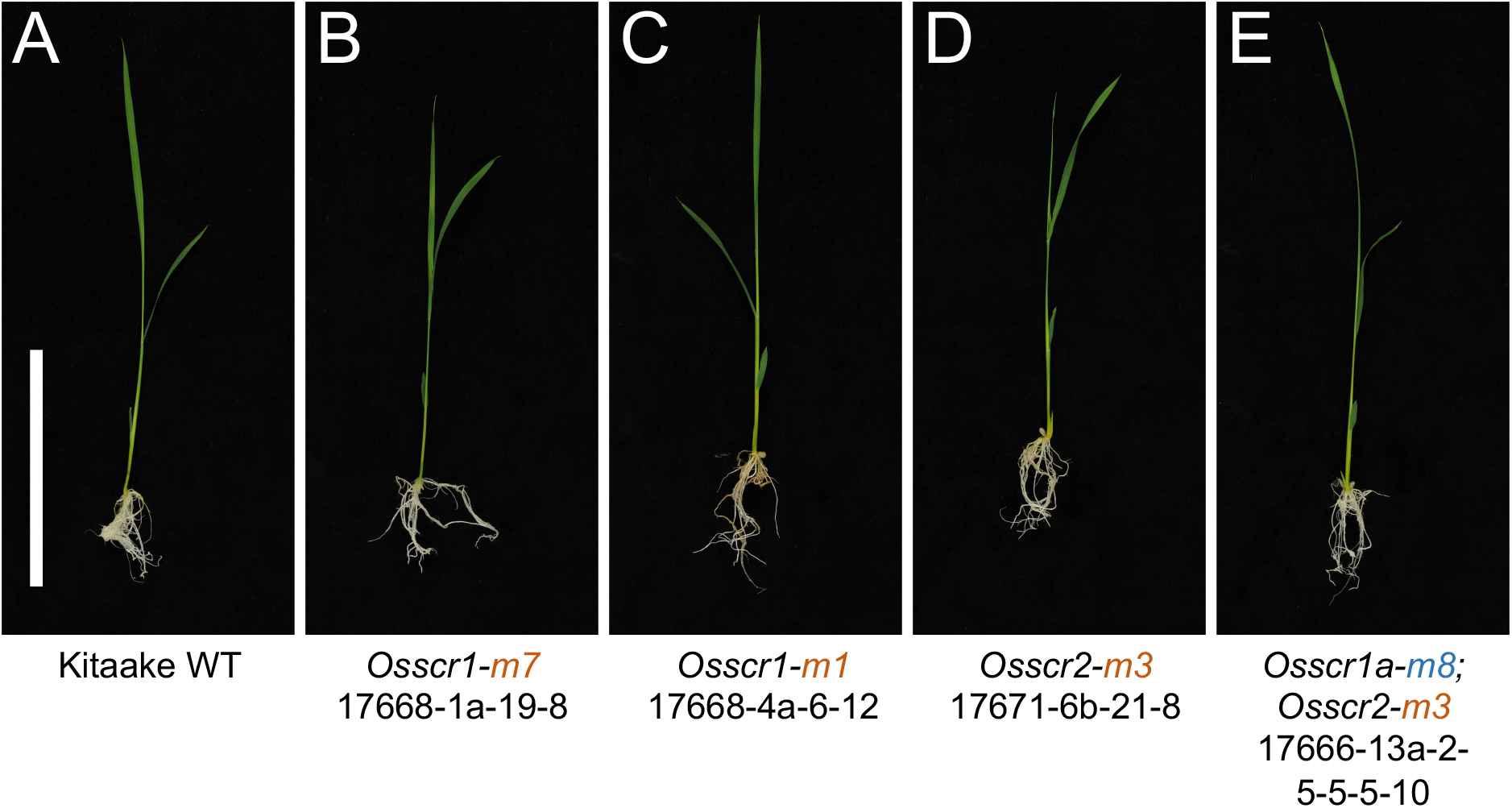
*Osscr1* and *Osscr2* single mutants grow normally. **A-E)** Photographs of WT (A), *Osscr1-m7* (B), *Osscr1-m1* (C), *Osscr2-m3* (D) and *Osscr1-m8;Osscr2-m3* (E) plants taken 13 days after germination. Orange alleles indicate out-of-frame mutations whereas blue alleles indicate an in-frame mutation not predicted to alter protein function. All mutants were obtained from independent lines or constructs, as indicated underneath the allele names. Scalebar in A is 10 cm and all images are at the same magnification.

**Figure S3.**
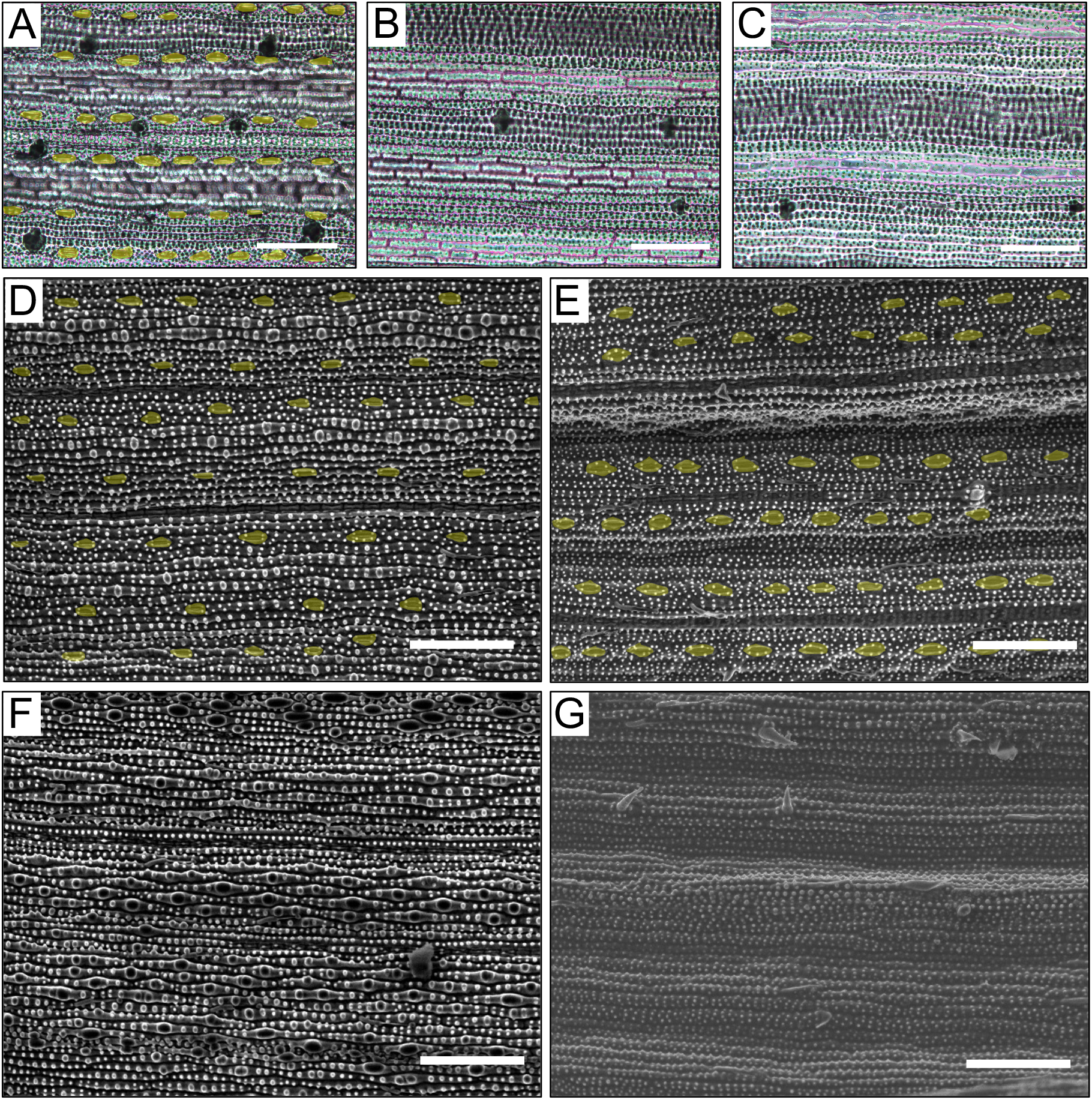
Stomatal phenotypes are consistent on both sides of the leaf and not an artefact caused by taking impressions. **A-C)** Adaxial impressions of WT (A), *Osscr1-m7;Osscr2-m3* (B) and *Osscr1-m7;Osscr2-m10* (C) leaf 5. Scalebars are 100μm. **D-G)** Scanning electron microscope images of leaf 5 for kitaake WT (D and E) or *Osscr1-m7;Osscr2-m3* (F and G) abaxial (D and F) and adaxial (E and G) surfaces. Scalebars are 100μm. Stomata are false coloured yellow.

**Figure S4.**
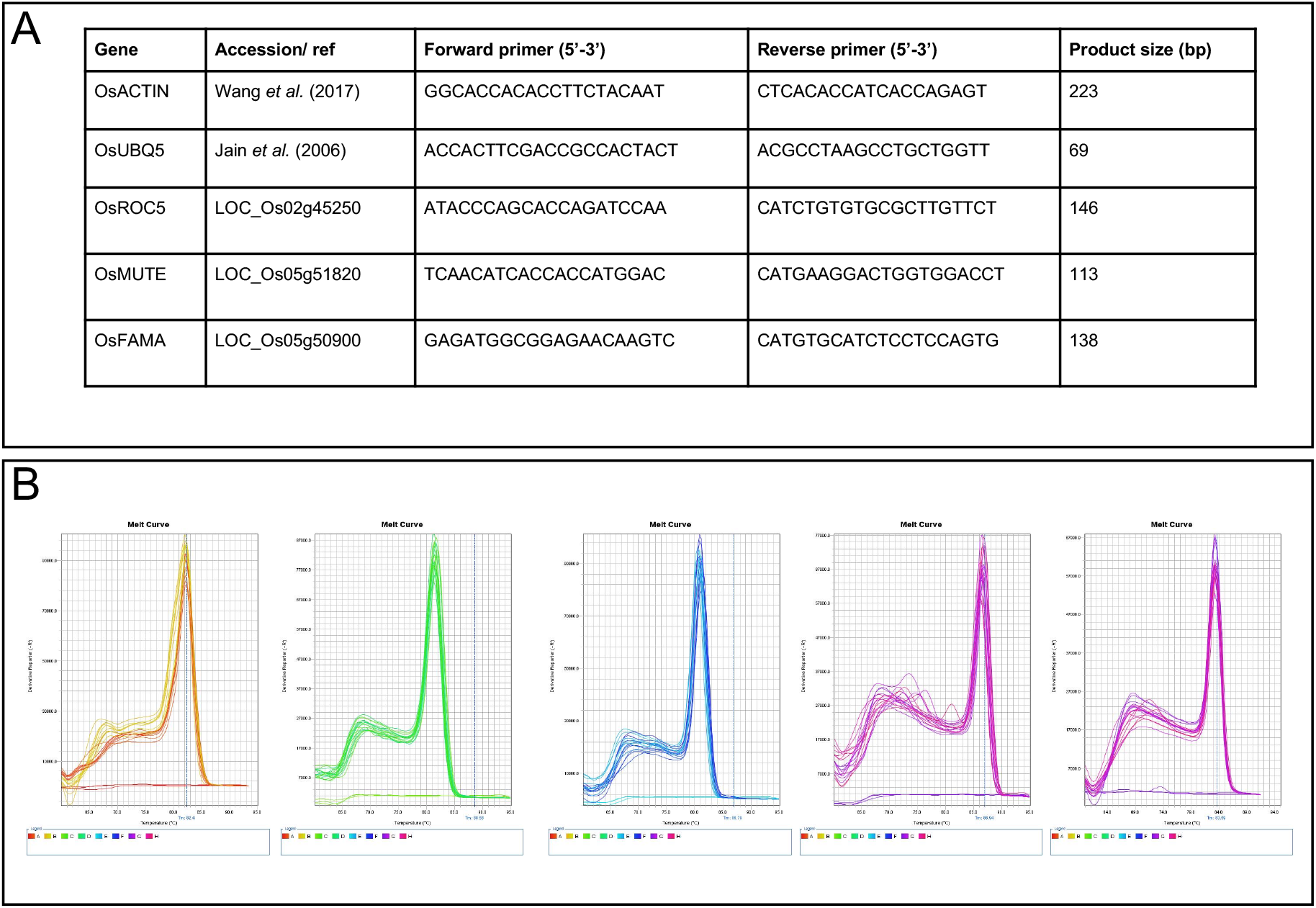
Quantitative RT-PCR primer design and validation. **A)** Primer sequences and product sizes. **B)** Melt-curves for each primer pair used in the study.

